# The mitochondrial Na^+^/Ca^2+^ exchanger, NCLX, mediates PDE2 dependent neuronal survival and learning

**DOI:** 10.1101/2021.07.08.451717

**Authors:** Maya Rozenfeld, Ivana Savic Azoulay, Tsipi Ben Kasus Nissim, Alexandra Stavsky, Michal Hershfinkel, Ora Kofman, Israel Sekler

## Abstract

Impaired phosphodiesterase (PDE) function and mitochondrial Ca^2+^ - [Ca^2+^]m signaling leads to cardiac failure, ischemic damage and dysfunctional learning and memory. Yet, a causative link between these pathways is unknown. Here, we fluorescently monitored [Ca^2+^]m transients in hippocampal neurons evoked by caffeine followed by depolarization. [Ca^2+^]m efflux was apparent in WT but diminished in neurons deficient in the mitochondrial Na^+^/Ca^2+^ exchanger NCLX. Surprisingly, neuronal depolarization-induced Ca^2+^ transients alone failed to evoke strong [Ca^2+^]m efflux in WT neurons. Caffeine is also a PDE inhibitor. Pretreatment with the PDE2 inhibitor Bay 60-7550 rescued [Ca^2+^]m efflux triggered by neuronal depolarization. Inhibition of PDE2 acted by diminishing the Ca^2+^ dependent reduction of mitochondrial cAMP, thereby promoting NCLX phosphorylation. Selective PDE2 inhibition also enhanced [Ca^2+^]m efflux triggered by neuromodulators. We found that protection of neurons against excitotoxic insults, conferred by PDE2 inhibition, was diminished in NCLX KO neurons, thus is NCLX dependent. Finally, administration of Bay 60-7550 enhanced new object recognition learning in WT but not in NCLX KO mice. Our results identify a long-sought link between PDE and [Ca^2+^]m signaling thereby providing new therapeutic targets.

## Introduction

Mitochondrial Ca^2+^ - [Ca^2+^]m signaling plays a double-edged role, with its uptake primarily meditated by MCU (Kirichok, Krapivinsky et al. 2004, Baughman, Perocchi et al. 2011), a specific Ca^2+^ channel that conducts [Ca^2+^]m permeation powered by the steep (∼ −180mV) mitochondrial membrane potential (Kirichok, Krapivinsky et al. 2004). Ca^2+^ is then pumped out of the mitochondria by an electrogenic mitochondrial Na^+^/Ca^2+^ exchanger NCLX (Palty, Silverman et al. 2010, De Stefani, Raffaello et al. 2011). The latter is much slower than MCU and therefore constitute a rate limiting step in the [Ca^2+^]m cycle (Traaseth, Elfering et al. 2004). The rise in [Ca^2+^]m activates several enzymes of the Krebs cycles (Rutter 1990, Traaseth, Elfering et al. 2004) and thereby the link between global Ca^2+^ rise and the energetic supply is required for many subsequent signaling and molecular events. By modulating the local Ca^2+^ regulation at the cell membrane and at the ER, mitochondria allosterically regulate the activity of receptors and ion channels (Poburko, Liao et al. 2009). Impaired balance between [Ca^2+^]m influx and efflux or change in [Ca^2+^]m buffering, however can lead to uncontrolled [Ca^2+^]m rise, the so-called [Ca^2+^]m overload, that triggers opening of the mitochondrial permeability pore (mPTP) and mitochondrial swelling (Rasola and Bernardi 2011). The second is an early hallmark event in ischemic and neurodegenerative syndromes. Remarkably the conditional KO of NCLX leads to profound [Ca^2+^]m overload and heart failure or Alzheimer’s disease (AD) symptoms in a mouse AD model (Luongo, Lambert et al. 2017, Jadiya, Kolmetzky et al. 2019). PKA dependent phosphorylation of NCLX upregulates [Ca^2+^]m efflux and promotes the survival of Parkinson disease associated PINK 1 deficient dopaminergic neurons (Kostic, Ludtmann et al. 2015, Kostic, Katoshevski et al. 2018). Still, the identity of key molecular players in the PKA pathway that control NCLX physiologically and thereby [Ca^2+^]m efflux (or failure thereof) in pathophysiological syndromes remain elusive.

Recent studies suggest that caffeine consumption lowers the risk of cardiovascular and neurodegenerative syndromes (Bhatti, O’Keefe et al. 2013). Caffeine is a potent albeit nonselective inhibitor of phosphodiesterase (PDEs), but how the health-related effects of caffeine are linked to PDEs is unknown. PDEs, by degrading cAMP, control the PKA pathway and are of major therapeutic importance (Technikova-Dobrova, Sardanelli et al. 2001, Acin-Perez, Salazar et al. 2009). For example, inhibitors of PDE5 are effective drugs for treating erectile dysfunction or pulmonary hypertension (Corbin 2004, Hemnes and Champion 2006) and PDE4 blockers are used to treated psoriasis and alcoholic addiction (Gooderham and Papp 2015, Logrip 2015). Among the PDEs, PDE2 is uniquely targeted also to the mitochondrial matrix and PDE2A is its only known expressed isoform of this enzyme (Acin-Perez, Russwurm et al. 2011). Numerous studies indicate that selective inhibitors of PDE2 protect against heart failure (Zaccolo and Movsesian 2007), promote neuronal survival and improve learning and memory in mice (Boess, Hendrix et al. 2004, Monterisi, Lobo et al. 2017). Still, how PDE2 controls these processes and whether it is linked to the mitochondria is poorly understood.

## Results

We first aimed to monitor mild versus strong [Ca^2+^]m signals triggered in WT and NCLX KO mouse primary hippocampal neurons preloaded with the [Ca^2+^]m sensor Rhod2-AM (we achieve spatially specific mitochondrial loading, as reported previously (Kostic, Ludtmann et al. 2015)). Consistent with previous studies (Walz, Baumann et al. 1995), application of caffeine triggered multiple small [Ca^2+^]m transients (Figures 1A-B). Subsequent application of high K^+^ Ringer’s buffer (5mM) induces neuronal depolarization and drives substantial Ca^2+^ entry that results in a much stronger [Ca^2+^]m response (Figures 1A). In NCLX KO neurons, the caffeine induced [Ca^2+^]m signal was less spike-like, leading to a consequent gradual and continuous Ca^2+^ accumulation in mitochondria (Figures 1A-C and 1E). Furthermore, while neuronal depolarization triggered a small influx phase followed by fast [Ca^2+^]m efflux in the WT neurons (Figures 1A, 1D and 1F), in NCLX KO neurons depolarization triggered higher [Ca^2+^]m influx and profound inhibition of [Ca^2+^]m efflux (Figure 1A, 1D and 1F). The depolarization-dependent [Ca^2+^]c response showed no significant differences between WT and NCLX KO neurons (Figures 1G-H), while cytosolic Ca^2+^ oscillations were still significantly different (Figure S1A). The results of this part indicate that caffeine-dependent spike-like signals and depolarization-dependent transient [Ca^2+^]m rises are controlled by NCLX in hippocampal neurons.

**Figure 1.**
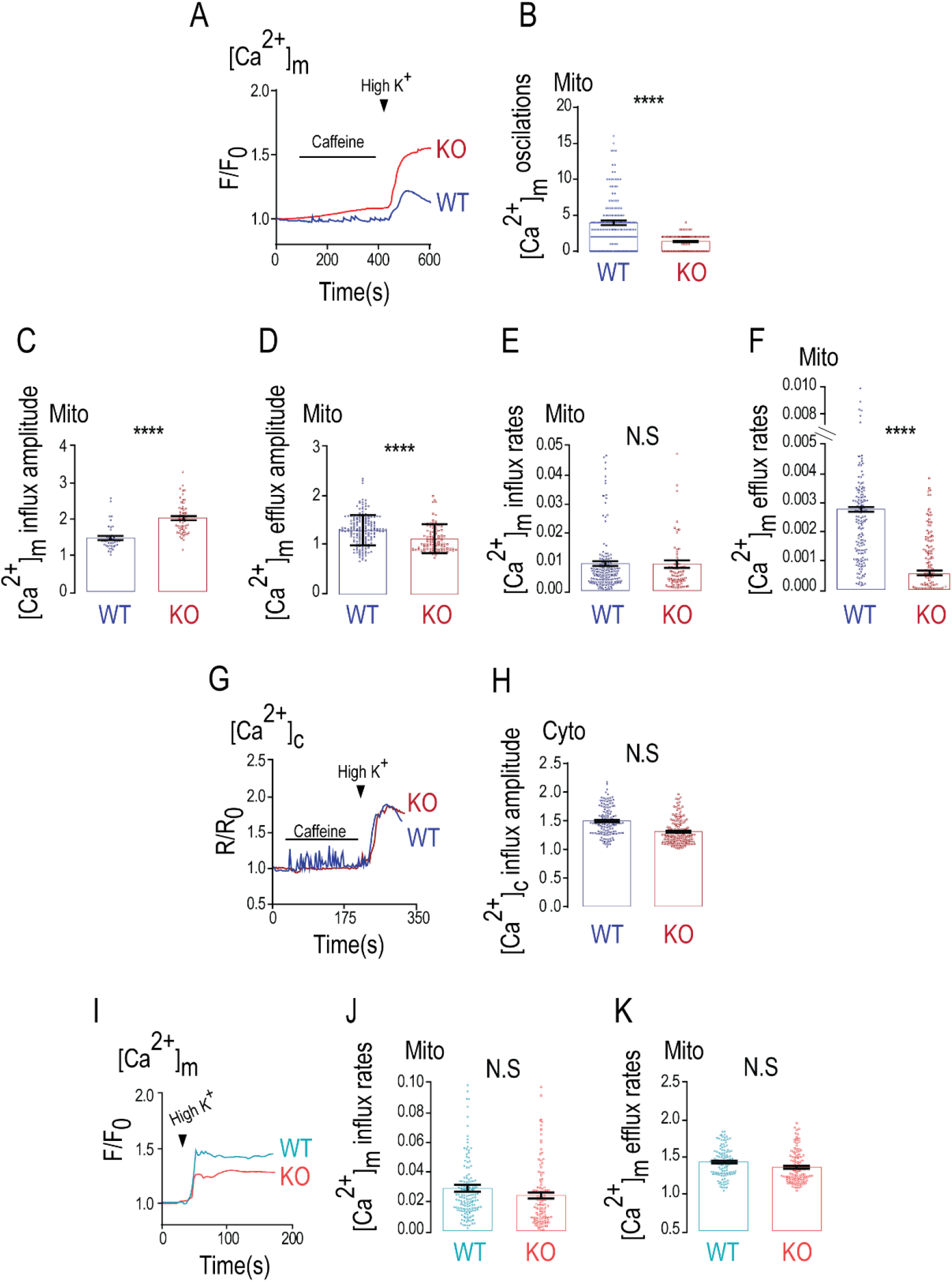
Mitochondrial Ca^2+^ extrusion is caffeine dependent in WT neurons, but not in NCLX KO neurons. (A) Representative fluorescence traces of mitochondrial Ca^2+^ [Ca^2+^]m changes monitored in WT and NLCX KO primary cultured hippocampal neurons (DIV 10-15). Neurons were preloaded with Rhod2-AM (1 µM) and initially superfused with Ringer’s solution at pH 7.4 to stabilize the base line. The [Ca^2+^]m signals were first triggered by caffeine (2mM) containing Ringer’s solution and then neuronal depolarization trigged by high K^+^ Ringer’s solution; (B) Quantification of [Ca^2+^]m oscillations showed in (A) for WT (n=162) and NCLX KO neurons (n=126); (C) Quantification of [Ca^2+^]m influx amplitude showed in (A) for WT (n=38) and NCLX KO neurons (n=54); (D) Quantification of [Ca^2+^]m efflux amplitude showed in (A) for WT (n=100) and NCLX KO neurons (n=115); (E) Quantification of [Ca^2+^]m influx rates showed in (A) for WT (n=38) and NCLX KO neurons (n=54); (F) Quantification of [Ca^2+^]m efflux rates showed in (A) for WT (n=164) and NCLX KO neurons (n=106); (G) Representative fluorescence traces of [Ca^2+^]c changes in WT and NCLX KO neurons preloaded with Fura 2-AM (1 µM) evoked by caffeine and high K^+^ Ringer’s solution as in (A); (H) Quantification of [Ca^2+^]c influx amplitude changes during the maximum phases of the data showed in (G) for WT (n=166) and NCLX KO neurons (n=99); (I) Representative fluorescence traces of [Ca^2+^]m changes monitored in WT and NLCX KO. [Ca^2+^]m signals were evoked only by high K^+^ Ringer’s solution; (J) Quantification of [Ca^2+^]m influx rates of data showed in (I) for WT (n=160) and NCLX KO neurons (n=114); (K) Quantification of [Ca^2+^]m efflux rates showed in (I) for WT (n=138) and NCLX KO neurons (n=145); All summary data represent mean ± SEM, **** p < 0.0001, N.S - non-significant.

We next sought to compare the [Ca^2+^]m response trigged only by the high K^+^ dependent depolarization (Figures 1I-K, S1B-C) in WT and NCLX KO neurons, reasoning that it will show the same gap in [Ca^2+^]m efflux between WT and NCLX KO neurons. Mitochondrial influx rates did not differ (Figure 1J), while mitochondrial influx amplitude was somewhat smaller in the NCLX KO neurons compared to WT (Figure 1SB), likely due to the loss of efflux pathway. Surprisingly, without the initial caffeine exposure, we no longer saw any difference between [Ca^2+^]m efflux rates in WT versus KO derived neurons following depolarization (Figures 1K and S1C), versus the large difference seen with caffeine pretreatment. We also show a smaller [Ca^2+^]c influx in the NCLX KO neurons (Figure S1D-E), consistent with greater mitochondrial uptake when main efflux pathway is knocked out. Thus, our results suggest that caffeine, independently of its Ca^2+^ ER release, had an additional effect of augmenting [Ca^2+^]m efflux in neurons via NCLX activation.

Caffeine has a well-documented regulatory effect on the PKA pathway and is known to inhibit PDEs (Ku, Lee et al. 2011, Zeitlin, Patel et al. 2011). Since our previous studies indicated that PKA plays a key role in regulating NCLX activity (Kostic, Ludtmann et al. 2015, Kostic, Katoshevski et al. 2018), we reasoned that the stimulating effect of caffeine on [Ca^2+^]m efflux by NCLX may be due to increased cAMP and therefore is PKA dependent. Hence, we first compared the [Ca^2+^]m responses triggered by caffeine followed by high K^+^ in control versus PKA inhibitor - H89 pretreated WT neurons (Figure 2A-D). Notably, PKA inhibition mimicked the NCLX KO (Figure 1A-B) in strong reduction of caffeine dependent [Ca^2+^]m spike-like activity, dramatic slowdown of [Ca^2+^]m efflux rate following neuronal depolarization (Figures 2A-B and 2D) and increase of the [Ca^2+^]m influx amplitude (Figure S2A), while not influencing influx rates (Figure 2C). In addition, and consistent with previous studies (Sang, Dick et al. 2016), the PKA inhibition also reduced peak [Ca^2+^]c response (Figure 2E-F). Taken together, our results suggest that caffeine is upregulating NCLX activity through the PKA pathway and that this regulation is critical for activation of [Ca^2+^]m efflux by this exchanger in neurons.

**Figure 2.**
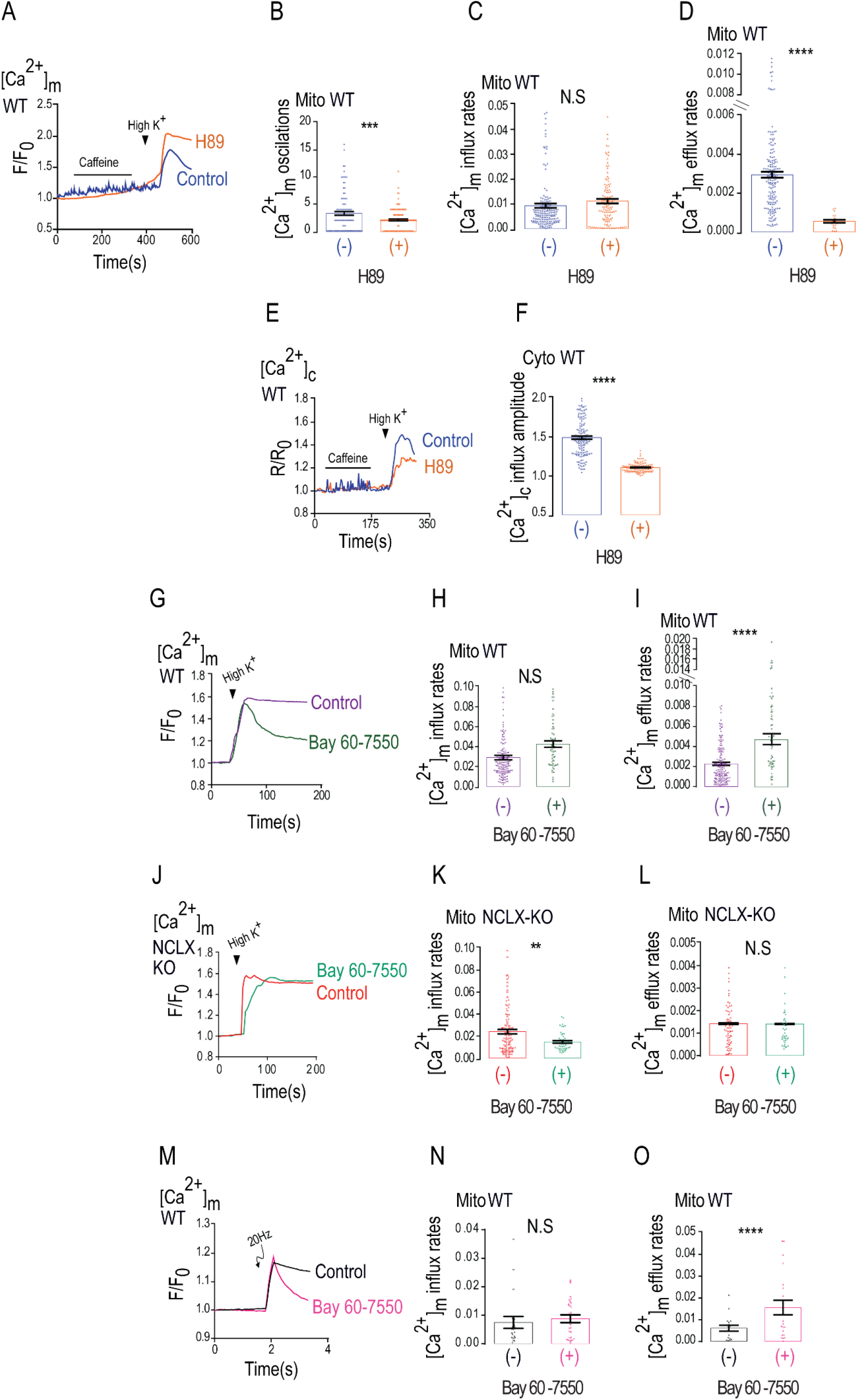
H89 inhibits Ca^2+^ efflux in WT neurons while PDE2 inhibitor mimics caffeine effect. (A) Representative fluorescence traces of [Ca^2+^]m changes monitored in control and H89 pretreated WT primary cultured hippocampal neurons. Neurons were loaded with Rhod2-AM (1 µM), initially superfused with caffeine followed by high K^+^ Ringer’s solution as in Figure 1A; (B) Quantification of [Ca^2+^]m oscillations showed in (A) for control (n=105) and H89 pretreated WT neurons (n=133); (C) Quantification of [Ca^2+^]m influx rates showed in (A) for WT (n=164) and H89 pretreated WT neurons (n=127); (D) Quantification of [Ca^2+^]m efflux rates showed in (A) for WT (n=164) and H89 pretreated WT neurons (n=30); (E) Representative fluorescence traces of [Ca^2+^]c changes in WT and H89 pretreated WT neurons done as in (Figure 1G); (F) Quantification of [Ca^2+^]c influx amplitude changes during the maximum phases of the data showed in (E) for control (n=138) and H89 pretreated WT neurons (n=124); (G) Representative fluorescence traces of [Ca^2+^]m changes monitored in control and Bay 60-7550 pretreated WT neurons. [Ca^2+^]m signals were evoked only by high K^+^ Ringer’s solution; (H) Quantification of [Ca^2+^]m influx rates changes showed in (G) for control (n=150) and Bay 60-7550 pretreated WT (n=60); (I) Quantification of [Ca^2+^]m efflux rates changes showed in (G) for control (n=155) and Bay 60-7550 pretreated WT (n=61); (J) Representative fluorescence traces of [Ca^2+^]m changes monitored in control NCLX KO and Bay 60-7550 pretreated NCLX KO neurons. [Ca^2+^]m signals were evoked only by high K^+^ Ringer’s solution; (K) Quantification of [Ca^2+^]m influx rates changes showed in (J) for control NCLX KO (n=107) and Bay 60-7550 pretreated NCLX KO neurons (n=52); (L) Quantification of [Ca^2+^]m efflux rates changes showed in (J) for control NCLX KO (n=90) and Bay 60-7550 pretreated NCLX KO neurons (n=50); (M) Representative fluorescence traces of [Ca^2+^]m changes monitored in control and Bay 60-7550 pretreated WT neurons. Neurons were preloaded with Rhod2-AM (1 µM), initially superfused with Ringer’s solution and then electrically stimulated by 5 consecutive pulses, each of 20 Hz frequency; (N) Quantification of [Ca^2+^]m influx rate changes during the maximum phases showed in (M) for WT (n=18) and Bay 60-7550 pretreated WT neurons (n=23); (O) Quantification of [Ca^2+^]m efflux rates showed in (M) for WT (n=19) and Bay 60-7550 pretreated WT neurons (n=24); All summary data represent mean ± SEM., ** p < 0.01, *** p < 0.001 **** p < 0.0001, N.S- non-significant.

Caffeine regulation of PKA is primarily mediated by inhibition of phosphodiesterase’s (Ribeiro and Sebastiao 2010), with the highest affinity to PDE2 (Technikova-Dobrova, Sardanelli et al. 2001, Acin-Perez, Salazar et al. 2009). Remarkably, PDE2 is the only PDE form that is targeted to the mitochondria matrix (Acin-Perez, Russwurm et al. 2011). We therefore asked if PDE2 is linked to NCLX regulation. We applied high K^+^ induced depolarization in the absence or presence of the selective PDE2 blocker Bay 60-7550. Addition of Bay 60-7550 rescued the depolarization dependent [Ca^2+^]m efflux, compared to control and similarly to caffeine (Figure 2G-I). No major effect on mitochondrial influx and a minor effect on [Ca^2+^]c responses were monitored (Figure 2C, S2B and S2D-E). In contrast to the strong effect of PDE2 inhibition on WT, in NCLX KO neurons the application of Bay 60-7550 had no effect on [Ca^2+^]m and [Ca^2+^]c responses (Figures 2J-L and S2C-E). This set of results suggest that PDE2 is regulating [Ca^2+^]m efflux by NCLX in neurons.

To further test the role of PDE2 using a depolarization-independent strategy, we monitored [Ca^2+^]m efflux in electrically stimulated (5 pulses, 20 Hz stimulation) neurons and found that Bay 60-7550 was similarly effective in rescuing the [Ca^2+^]m efflux in such stimulated neurons, while not changing the [Ca^2+^]m influx (Figure 2M-O). This set of experiments indicate that inhibition of PDE2 is required for activation of [Ca^2+^]m efflux by NCLX in neurons.

We next ask if PDE2 activity, invoked by the [Ca^2+^]m transients, is regulating mitochondrial cAMP content. To monitor mitochondrial cAMP, we targeted the cAMP sensor Pink Flamindo (PF) (Harada, Ito et al. 2017) to the mitochondria by adding a mitochondrial targeting domain (See Materials and methods section 4) (Figure 3A) and found that the reconfiguration of this reporter led to its mitochondrial targeting (Figure 3B). To determine if Bay 60-7550 has an effect on mitochondrial cAMP levels we fluorescently monitored mitochondrial cAMP in neuronal SH-SY5Y cells and primary hippocampal neurons expressing this mitochondrial targeted PF (mtPF) by triggering ATP dependent purinergic or depolarization dependent (high K^+^) Ca^2+^ signals respectively (Figure 3C-F). In SH-SY5Y cells ATP evoked a rise in mitochondrial cAMP (Figure 3C-D). In contrast, and consistent with previous studies (Bubb, Trinder et al. 2014), SH-SY5Y cells pretreated with Bay-60-7550 showed a larger rise in mitochondrial cAMP. In primary neurons, depolarization triggered a fast drop in mitochondrial cAMP that was largely suppressed in Bay 60-7550 pretreated neurons (Figure 3E-F).

**Figure 3.**
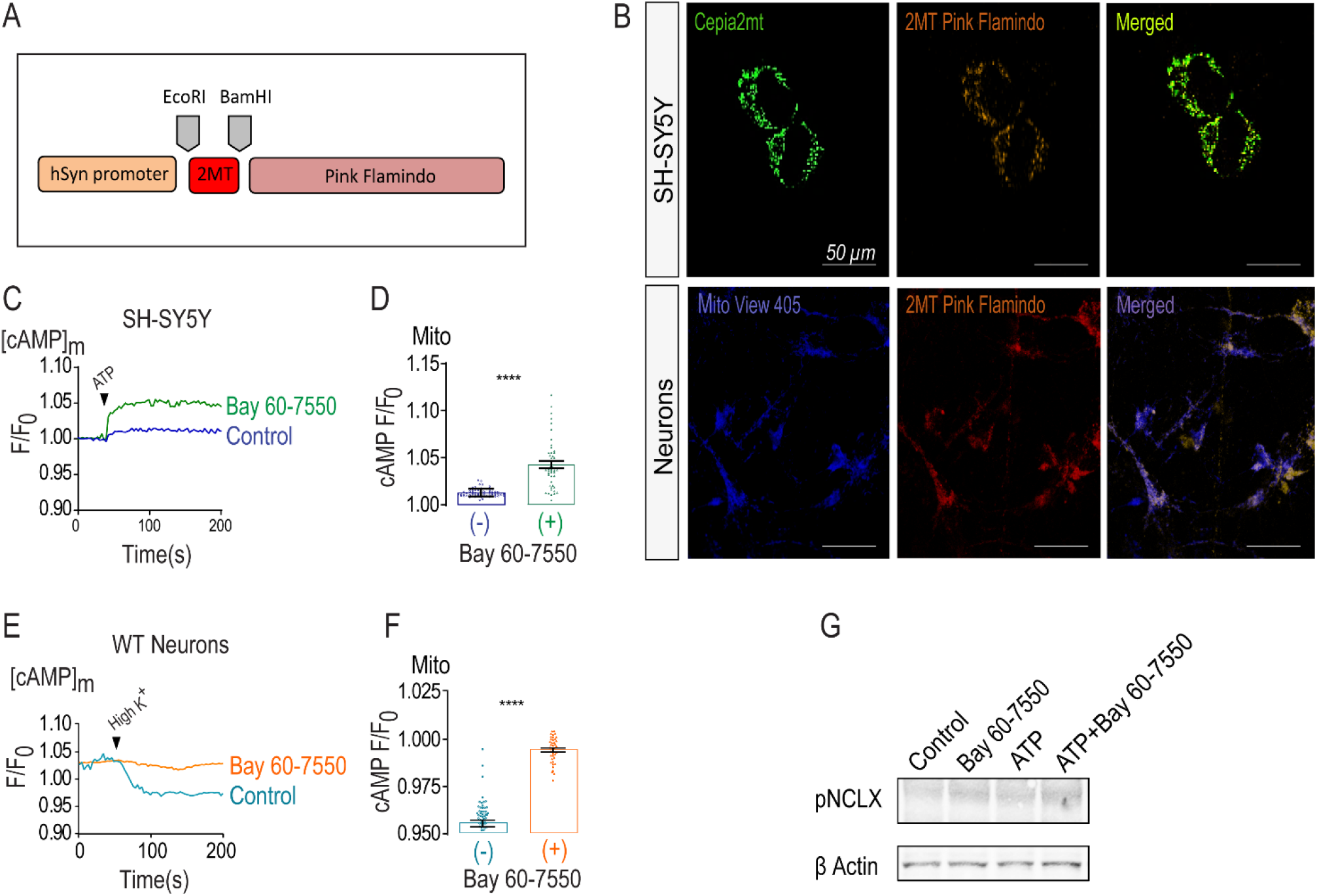
Bay 60-7550 increases NCLX activity by stimulating its phosphorylation. (A) Scheme of the mitochondrial Pink Flamindo construct; (B) Representative confocal images of SH-SY5Y expressing the [Ca^2+^]m reporter CEPIA 2-AM (1µg)-green, Pink Flamindo AAV (mtPF)- red and their mitochondrial colocalization -yellow, scale bar indicates 50µm (upper); Representative confocal images of primary hippocampal neurons expressing the [Ca^2+^]m reporter Mito-View 405 (100nM) -blue, mtPF- red and their mitochondrial colocalization- purple, scale bar indicates 50µm (lower); (C) Representative fluorescence traces of mitochondrial cAMP changes in control and Bay 60-7550 pretreated SH-SY5Y cells expressing mtPF. Cells were superfused with Ringer’s solution followed by addition of ATP (100 µM) to evoke a Ca^2+^ response; (D) Quantification of cAMP changes showed in (D) of control (n=71) and Bay 60-7550 treated SH- SY5Y cells (n=46); (E) Representative fluorescence traces of cAMP changes in WT and Bay 60-7550 pretreated WT primary hippocampal neurons previously expressing the mtPF. Neurons were superfused with Ringer’s solution followed by high K^+^ to trigger a Ca^2+^ transient; (F) Quantification of cAMP changes showed in (F) for control (n=59) and Bay 60-7550 pretreated WT neurons (n=40); (G) Immunoblot of SH-SY5Y cells, reacted with anti pNCLX Ab, showing the NCLX phosphorylation at its PKA dependent S258 NCLX phosphorylation site. SH-SY5Y cells were pretreated either with Bay 60-7550 (1µM, 1 hour), ATP (100µM, 1 min) or both. Actin was used as loading control. All summary data represent mean ± SEM., **** p < 0.0001.

We then asked if the mitochondrial NCLX phosphorylation, at its S258 PKA phosphorylation site, is the target of PDE2. Here we focused on the SH-SY5Y cells because of the restricted selectivity of the pS258 antibody to human NCLX (Kostic, Katoshevski et al. 2018) (See Materials and methods, section 8). We found that the inhibition of PDE2 by Bay 60-7550 enhanced NCLX phosphorylation at the S258 PKA dependent position (Figure 3G). Our results indicate that the activation of NCLX by the PDE2 inhibition is mediated by rise in mitochondrial cAMP by triggering NCLX phosphorylation at its PKA S258 phosphorylation site.

We next interrogated the role of PDE2 in controlling the [Ca^2+^]m transients evoked by the neuromodulators Gq coupled vasopressin (VP) and Gs coupled norepinephrine (NE). Their application evoked cytosolic Ca^2+^ transients (Figure S3) followed by mitochondrial influx but not substantial efflux of Ca^2+^ (Figure 4). In contrast, pretreatment of WT neurons with Bay-60-7550 revealed VP and NE dependent [Ca^2+^]m efflux in WT but not in NCLX KO neurons (Figures 4A-D and 4E-H), indicating that both NCLX and inhibition of PDE2 were required to activate neurotransmitters dependent [Ca^2+^]m efflux. [Ca^2+^]c transients mimicked the results shown for [Ca^2+^]m (Figure S3A-H). Interestingly co-application of VP and NE was sufficient to reveal [Ca^2+^]m efflux (Figures 4I, 4K and S3I-L). When Bay 60-70550 co-applied with NE and VP it strongly amplified [Ca^2+^]m efflux resulting in a large drop in [Ca^2+^]m that was well below resting set point in WT neurons (Figure 4I-J). No efflux was monitored in NCLX KO neurons in the presence or absence of Bay-60-7550 (Figure 4K-L). We suggest that inhibition of PDE2 is required to reconstitute the neurotransmitters dependent [Ca^2+^]m efflux. Furthermore, NE and VP play a synergistic role in amplifying [Ca^2+^]m efflux that is fully unmasked upon PDE2 inhibition.

**Figure 4.**
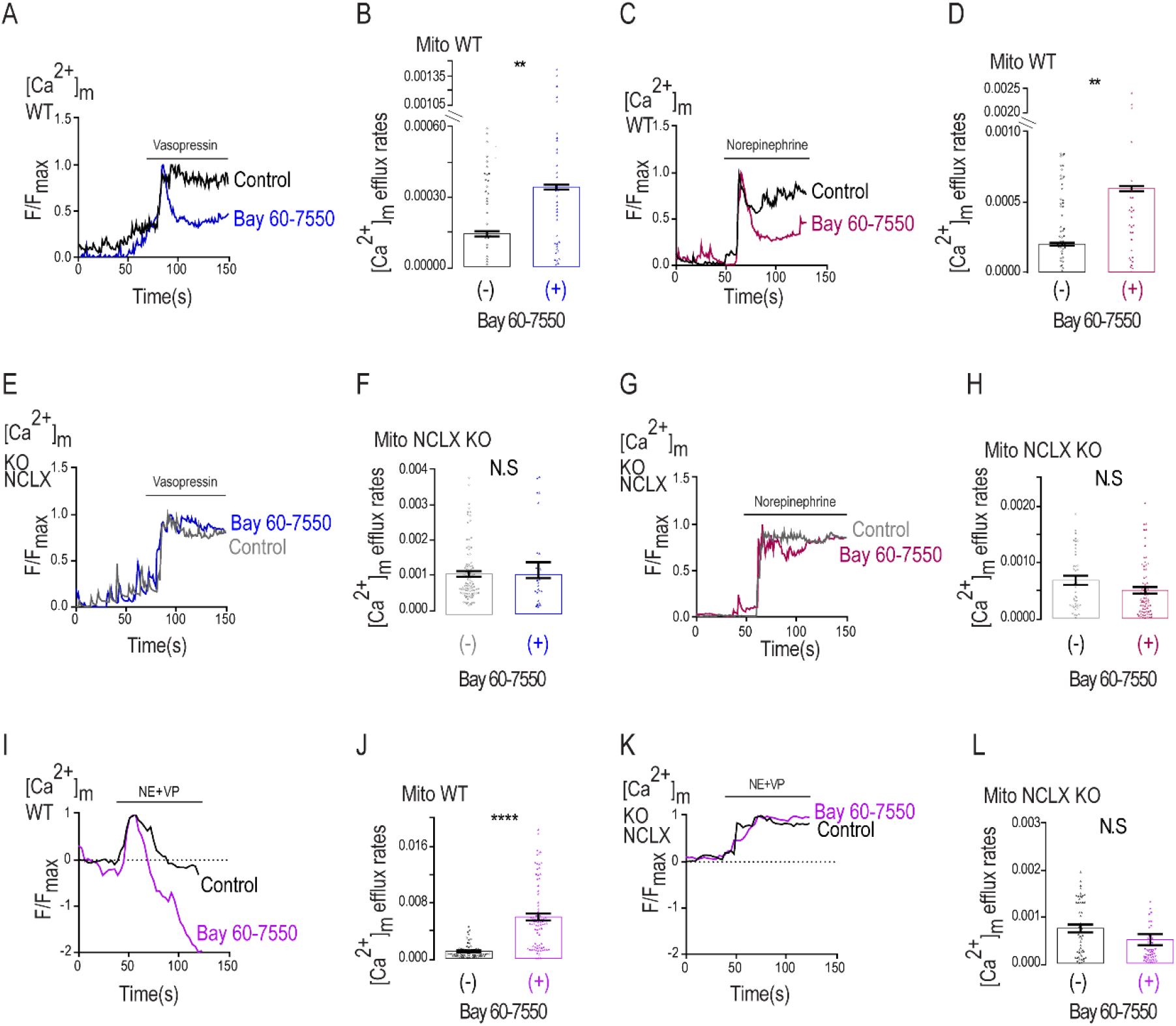
VP and NE trigger mitochondrial Ca^2+^ influx followed by Bay 60-7550 enhanced NCLX mediated Ca^2+^ efflux. (A) Representative fluorescence traces of [Ca^2+^]m changes monitored in VP treated control and Bay 60-7550 pretreated WT neurons. Neurons were preloaded with Rhod2-AM, initially superfused with Ringer’s solution. (1 µM) VP (1 µM) containing Ringer’s solution was added as indicated by the bar; (B) Quantification of [Ca^2+^]m efflux rates showed in (A) for VP treated control (n=53) and Bay 60-7550 pretreated WT neurons (n=41); (C) Representative fluorescence traces of [Ca^2+^]m changes monitored in NE (5 µM) treated control and Bay 60-7550 pretreated WT neurons as in (A); (D) Quantification of [Ca^2+^]m efflux rates showed in (C) for control (n=62) and Bay 60-7550 pretreated WT neurons (n=33); (E) Representative fluorescence traces of [Ca^2+^]m changes in VP treated control NCLX KO and Bay 60-7550 pretreated NCLX KO neurons using the same experiment paradigm shown in (A); (F) Quantification of [Ca^2+^]m efflux rates showed in (E) for VP treated control KO (n=70) and Bay 60-7550 pretreated NCLX KO neurons (n=30); (G) Representative fluorescence traces of [Ca^2+^]m changes NE treated control NCLX KO and Bay 60-7550 pretreated NCLX KO neurons as in (E); (H) Quantification of [Ca^2+^]m efflux rates showed in (G) for NE treated control NCLX KO (n=51) and Bay 60-7550 pretreated NCLX KO neurons (n=76); (I) Representative fluorescence traces of [Ca^2+^]m changes monitored of WT neurons treated with VP and NE co-applied at time interval indicated by the time bar (100s) in the absence or presence of Bay 60-7550 pretreated WT neurons as described in A; (J) Quantification of [Ca^2+^]m efflux rates showed in (I) for control (n=67) and Bay 60-7550 pretreated WT neurons (n=90); (K) Representative fluorescence traces of Ca^2+^]m changes monitored in NCLX-KO neurons treated with VP and NE of control NCLX KO and Bay 60-7550 pretreated NCLX KO neurons of the experiment performed as in (I); (L) Quantification of [Ca^2+^]m efflux rates showed in (K) for NCLX KO (n=50) and Bay 60-7550 pretreated NCLX KO neurons (n=52); All summary data represent mean ± SEM., ** p < 0.01, **** p < 0.0001, N.S- non-significant.

Inhibition of PDE has emerged as promising strategy in preventing neuronal death (Ding, Zhang et al. 2014). Considering the strong link of [Ca^2+^]m overload and neuronal survival, we asked if the reactivation effect of PDE2 blocker on [Ca^2+^]m efflux by NCLX will rescue neurons subject to an excitotoxic insult. We subjected cultured neurons to such an insult by pretreating them overnight with the glutamate uptake transport inhibitor-TBOA (Sharifullina and Nistri 2006, Corsini, Tortora et al. 2016, Yeh, Bulas et al. 2017). As shown previously (Soares, Meyer et al. 2017), Bay 60-7550 was effective in preventing the TBOA dependent neuronal death (Figure 5A-C). Remarkably the rescuing effect of Bay 60-7550 on neuronal survival was largely eliminated in NCLX KO neurons indicating that the PDE2 blocker is enhancing survival by acting through the activation of NCLX.

**Figure 5.**
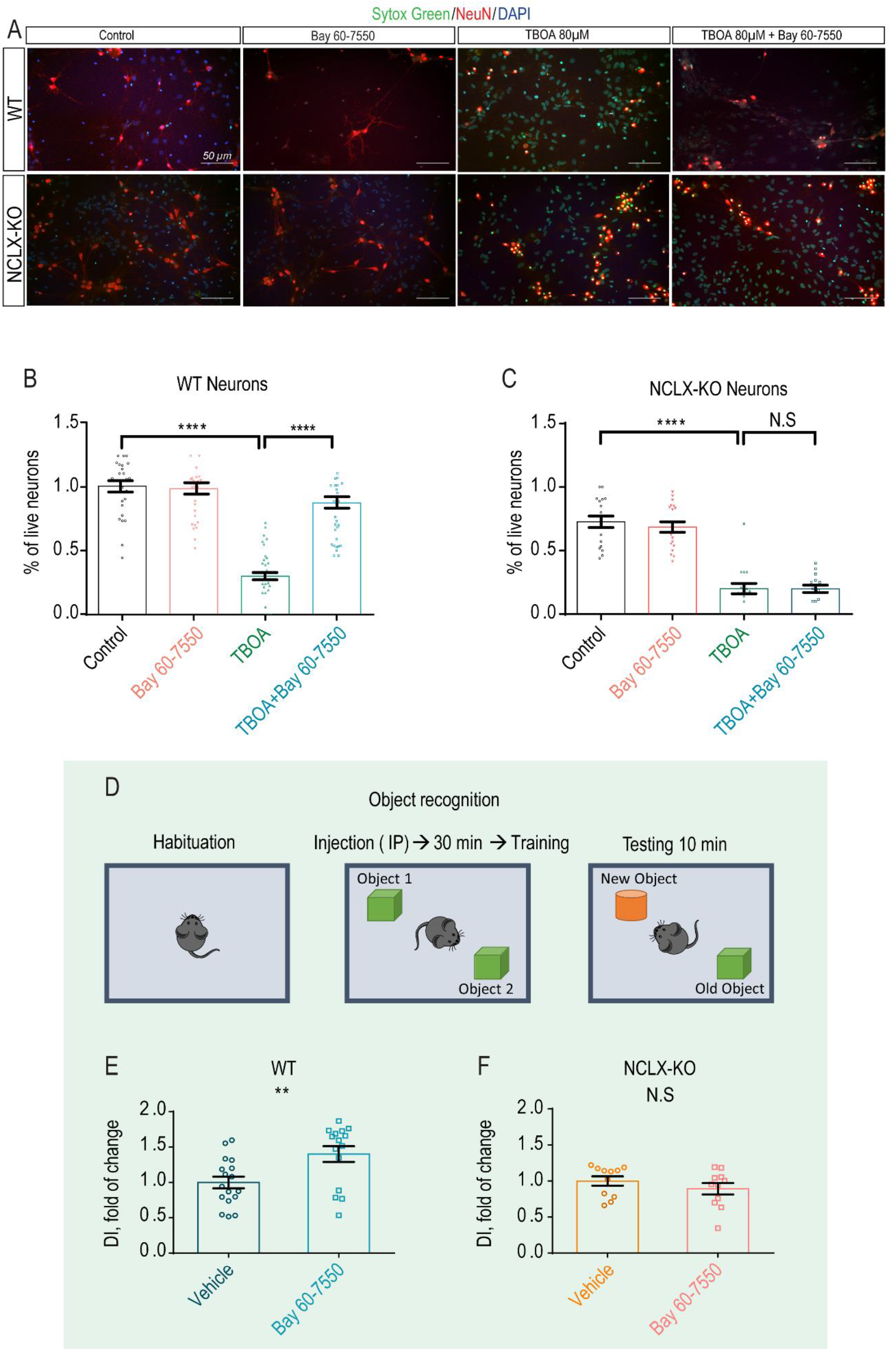
Bay 60-7550 promotes neuronal survival and mouse learning via NCLX regulation. (A) Representative images of WT and NCLX KO primary hippocampal neurons stained with Sytox Green, NeuN and DAPI for cellular death, specific neuronal and nuclei labeling, presented as control, Bay 60-7550 (1 µM, 1-hour pretreatment), TBOA (80 µM, overnight pretreatment) and TBOA+Bay 60-7550 (overnight and 1-hour pretreatment, respectively), scale bar indicates 50µm; (B) Quantification of live WT neurons showed in (A) presented as percentage (n=43); (C) Quantification of live NCLX KO neurons showed in (A) presented as percentage (n=20); (D) Schematic illustration of the Bay 60-7550 treatment/object recognition experimental design; (E) Discrimination index of vehicle (control) vs Bay 60-7550 injected WT mice (n=15/17); (F) Discrimination index of vehicle (control) vs Bay 60-7550 NCLX-KO mice (n=11); All summary data represent mean ± SEM., ** p < 0.01, **** p < 0.0001, N.S- non-significant.

Finally, numerous studies show that inhibition of PDE2 is effective in enhancing learning and memory (Boess, Hendrix et al. 2004, Lueptow, Zhan et al. 2016), but the molecular link that couples PDE2 to these processes is unknown. Considering the role of mitochondria in controlling neuronal plasticity (Mattson 2007), we reasoned that the well-established effect of PDE2 inhibition on enhancing learning and memory may be linked to [Ca^2+^]m extrusion rescue via NCLX. To address this hypothesis, we subjected control and Bay 60-7050 injected WT and NCLX KO mice to a new object recognition test (Figure 5D-F). Bay 60-7550 injection enhanced discrimination between two objects in WT mice as previously reported, while it showed no significant effect of the same parameter in NCLX KO mice. Altogether these results indicate that NCLX is required for PDE2-dependent enhancement of new object recognition.

## Discussion

We studied the effect of caffeine on ER Ca^2+^ release in hippocampal neurons. To our surprise we found that caffeine also had a profound and lasting effect in activating [Ca^2+^]m efflux in these neurons. The PDE inhibitory activity of caffeine (Ribeiro and Sebastiao 2010) and its regulation of NCLX, led us to hypothesize that the regulation of [Ca^2+^]m efflux was mediated by PDE dependent NCLX. Notably, among PDEs, PDE2 has the highest affinity to caffeine. It is only PDE2 that is targeted to the mitochondria (Acin-Perez, Salazar et al. 2009), we therefore reasoned that PDE2 is regulating NCLX in neurons and our hypothesis is supported by the following findings: 1) Caffeine upregulation of [Ca^2+^]m efflux is reproduced by the selective PDE2 inhibitor Bay 60-7550 (Figure 2); 2) The upregulation of [Ca^2+^]m efflux by Bay 60-7550 is blunted in neurons derived from NCLX-KO neurons, indicating that PDE2 is controlling NCLX-mediated [Ca^2+^]m efflux (Figure 2 and 4); 3) Application of Bay 60-7550 enhances NCLX phosphorylation at its NCLX S258 site indicating that PDE2 is acting through the PKA phosphorylation site of NCLX (Figure 3); 4) Remarkably, the KO of NCLX eliminates hallmark physiological implications of PDE2 inhibition by enhancing neuronal survival and cognitive function (Figure 5).

The unique distribution of PDE2 in brain regions associated to cognitive function indicates that the enzyme is linked to these processes. Indeed, numerous studies employing PDE2 inhibitors show that inhibition of PDE2 enhances (Gomez and Breitenbucher 2013) cognitive function in AD model during regular learning paradigm and in particular new object recognition were also clinically tested (Boess, Hendrix et al. 2004, Lueptow, Zhan et al. 2016). Despite the intense interest in the neuronal PDE2 role, a major unsolved question that impedes the progress in the field is the identity of its cellular targets, which further mediates its neuronal effects. As described previously, we find that inhibition of PDE2 by Bay 60-7550 rescues cells against excitotoxic insults. The pro-survival effect of the PDE2 blocker is fully diminished in NCLX-KO neurons indicating that this pathophysiological aspect of PDE2 is mediated by controlling NCLX. Remarkably the well documented enhancement of new object recognition triggered by PDE2 blocker is also NCLX dependent. These results indicate that NCLX is the long-sought mitochondrial target for PDE2.

Although we don’t fully know how NCLX is linked to cognitive function, recent studies indicate that mitochondria are critical for providing the metabolic energy and anabolic production of molecules required for de-novo protein synthesis (Weinberg, Sena et al. 2015). Such proteins are essential for synaptic plasticity linked to the mentioned processes. Our results suggest that the temporal control of [Ca^2+^]m transients plays a role in facilitating this process which may be governed by NCLX. Of particular interest is our observation that [Ca^2+^]m response is temporally modulated by neuromodulators such as the G_s_ coupled NE and G_q_ coupled VP (Figure 4). Additionally, by facilitating effects of PDE2 on mitochondrial fluxes, NE and VP modulate PDE2 regulation on NCLX. Considering the major metabolic and anabolic cost in providing the energy for neuronal transmission and for plasticity-linked new protein translation, the key role of Ca^2+^ in tuning mitochondrial activity constitutes the link between PDE2 and PKA regulation of NCLX and may play role in controlling such processes. Interestingly, we recently found that a human genetic mutation of NCLX, impairing its Ca^2+^ transport activity, is associated with severe mental retardation (Stavsky, Stoler et al. 2021).

The brain may not be the only organ where the interaction between PDE2 and NCLX have pathophysiological importance. The conditional knock down of cardiac NCLX leads to lethal heart failure (Jadiya, Kolmetzky et al. 2019). Remarkably the inhibition of PDE2 effectively prevents heart failure in a murine model (Vettel, Lindner et al. 2017). Considering the link between PDE2 and NCLX, we suggest that the cardio-protective effect of PDE2 may be mediated via NCLX. We therefore predict that PDE2 inhibition may not protect against heart failure in cardiac NCLX-KO mice and that studies focusing on the PDE2-NCLX axis may provide new therapeutic tools for major cardiac and brain related pathologies.

To conclude, we show that in hippocampal neurons [Ca^2+^]m efflux is tonically suppressed but can be activated by caffeine. The strong effect of PDE2 inhibition on [Ca^2+^]m efflux by attenuation of mitochondrial cAMP degradation is linked to NCLX phosphorylation at its PKA phosphorylation site. Additionally, we show that the PDE2 block is critical for enhancing neuronal survival following excitotoxic insult only when NCLX is active. Finally, we suggest that the enhancement of learning and memory functions mediated by PDE2 inhibition are NCLX dependent.

## Materials and methods

### 1. Primary neuronal culture

C57BL6 Wild-Type (WT) (Envigo, C57BL Cat #6JRCCH5B043) 0-3 days old mouse pups were maintained as previously described (Gitler, Takagishi et al. 2004). NCLX KO mice were obtained from Jackson Laboratories (Cat #026242). This strain was generated at the Jackson Laboratory by injecting Cas9 RNA and an 18-mer guide sequence ATACTGGAGACGGCGTCT, which resulted in a 13 bp deletion (AGACGGCGTCTGG) in exon 2 beginning at chromosome 5 positive strand position 120, 513, 241 bp (GRCm38), which is predicted to cause amino acid sequence changes after residue 21 and an early truncation 33 amino acids late.

Animals were treated with the approvals and in accordance with the guidelines of the Ben-Gurion University Institutional Committee for Ethical Care and Use of Animals (Reference Number: IL 07-02-2019C).

Briefly, mice were housed in specific pathogen free facilities. Rodent care practices were under sterile conditions, with sterile supplies. Rodents Hosing conditions are: 12:12 light:dark cycles at 20-24°C and 30–70% relative humidity. Animals are free fed autoclaved rodent chow and have free access to reverse osmosis filtered water. Rodents are housed in individually ventilated GM500 (Tecniplast, Italy) cages in groups of maximum 5 mice per cage.

Primary culture of hippocampal neurons was done for each independent experiment from mouse pups as previously described (Gitler, Xu et al. 2004). Each mouse hippocampus was used to generate culture on 6 coverslips with 50,000 - 70,000 cells. Cultures were typically maintained at 37°C in a 5% CO_2_ humidified incubator for 10-15 days before experimentation. In fluorescent imaging related experiments neurons were easily distinguishable from glia: they appeared phase bright, had smooth rounded somata and distinct processes and laid just above the focal plane of the glial layer.

### 2. Genotyping

NCLX +/+ (WT) or NCLX -/- (KO) mice were used after genotyping using PCR of DNA isolated from mouse tail biopsy samples. Primers 5’- TGGCTCTGATACTGGAGACG-3’ and 5’- CATGGCAGTCTGGTTGACAC- 3’ amplified a 128 bp band from the wild-type allele whereas primers 5’ -TGCTCTGGGCTCCTGTCTTC- 3’ and 5’- CATGGCAGTCTGGTTGACAC- 3’ amplified a 115-bp band from the knockout allele. PCR products were run on 4% Nusieve agarose gel (Cat #50090, LONZA) for higher resolution of bands separation results (Witt, Jaruzelska et al. 1993).

### 3. Cell cultures and transfection

SH-SY5Y cell lines were cultured (37^0^C, 5% CO_2_) in Dulbecco’s modified Eagle’s medium (DMEM) supplemented with 10% fetal bovine serum (FBS), 2mM L-glutamine and 1% penicillin/streptomycin, as previously described (Palty, Ohana et al. 2004). Transfection of SH-SY5Y cells was performed using the calcium phosphate precipitation protocol as previously described (Palty, Ohana et al. 2004).

### 4. Generating 2mtPink Flamindo cAMP biosensor

Pink Flamindo vector, a biosensor for cAMP in mammalian cells was purchased from Addgene (Harada, Ito et al. 2017) (Cat #102356). In order to direct this sensor into the mitochondria, the 2MT signaling peptide was genetically introduced into Pink Flamindo vector using the 2MTGCaMP6m vector gifted from Diego De Stefani (Patron, Checchetto et al. 2014). A vector containing the cAMP biosensor together with the 2MT segment was constructed as follows: The 2MT segment was subcloned by PCR amplification of the 2MTGCaMP6m vector utilizing the following primers: forward, ATGCGAATTCCACCATGTCCGTCCTGAG and reverse GCTAGGATCCAGAACCAAGCTTCCCCTCCG. The PCR product (237 bp) was then ligated into the Pink Flamindo vector between the ECOR1 and BamH1 sites to generate the 2mtPink Flamindo.

### 5. 2mtPink Flamindo virus preparation

Viral particles were prepared as described previously (Tevet and Gitler 2016). Briefly, cDNAs of interest (2MTFlamindo and 2MTGCaMP6m) were subcloned by restriction/ligation into a plasmid containing adeno-associated virus 2 (AAV2) inverted terminal repeats flanking a cassette consisting of the the 0.47 kb human synapsin 1 promoter (hSyn) (Kugler, Kilic et al. 2003), the woodchuck post-transcriptional regulatory element and the bovine growth hormone polyA signal. Viral particles were produced in HEK-293 cells using both the pD1 and pD2 helper plasmids (Groh, de Kock et al. 2008), which encode the rep/cap proteins of AAV1 and AAV2, respectively. Primary cultures of hippocampal neurons were infected at DIV 5 and incubated for at least 7 days before imaging. Virion titer was individually adjusted to produce 75–90% infection efficiency.

To test 2mtPink Flamindo mitochondrial localization in SH-SY5Y cells, we used mitochondrial marker CEPIA2-mt (Addgene plasmid, Cat #58218) as previously described (Kostic, Ludtmann et al. 2015). In neurons Pink Flamindo was co-localized with Mito-View 405 (Biotium, Cat #70070-T). Live cells confocal imaging was performed on confocal microscope described in methods section 6.

### 6. Live fluorescence imaging

Experiments were done on imaging system consisted of Olympus IX73 inverted microscope and Cellsense division 1 Olympus software (Wendenstrasse, Hamburg, Germany) CooLed 4000 LED monochromator, and Q imaging cooled Retiga 6000 camera (Surrey, British Columbia, Canada). We used magnification 10x for cytosolic and 20x for mitochondrial measurements. For live cytosolic Ca^2+^ (Fura2-AM) imaging we used (Zeiss), Polychrome V monochromator (TILL Photonics, Planegg, Germany) and a SensiCam cooled charge-coupled device (PCO, Kelheim, Germany) as previously described (Kostic, Ludtmann et al. 2015).

Experiments done on WT, treated WT and NCLX KO live neurons stimulated electrically, were performed using the imaging system consisted of Nikon TiE inverted microscope driven by the NIS-elements software package (Nikon). The microscope was equipped with an Andor sCMOS camera (Oxford Instruments), a 20x XX NA and 40x 0.75 NA Super Fluor objectives (Nikon), EGFP and Cy3 TE-series optical filter sets (Chroma) as well as a BFP filter set (Semrock) as previously described (Stavsky, Stoler et al. 2021). The baseline fluorescence intensity of MitoGcaMP6m (F0) in each area of interest was the calculated average value measured in 20 successive images acquired before stimulation. The change in fluorescence (ΔF) at time (t) was calculated as F(t)-F0. A mean trace was determined for each experiment and then a grand average was calculated for each condition.

Live mitochondria co-localizations and whole fixed neurons images were taken on a NIKON C2Plus laser unit docked to a Nikon Eclipse Ti unit of the confocal microscope by using a 20X and 60X oil immersion objective respectively, as described (Leyton-Jaimes, Kahn et al. 2019).

In all live fluorescent experiments, cells were pre-loaded with the indicated ion specific fluorescent dye at the indicated concentrations for 30 min at room temperature using Ringer’s solution containing (in mM): 126 NaCl, 5.4 KCl (Sigma-Aldrich, Cat #P9333), 0.8 MgCl_2_, 20 HEPES, 1.8 CaCl_2_, 15 glucose (Gerbu, Cat #2028), with pH adjusted to 7.4 with NaOH (Sigma-Aldrich, Cat #221465) and supplemented with 0.1% bovine serum albumin (BSA, VWR, Cat #0332). After loading, cells were thoroughly washed in fresh dye-free Ringer’s solution for 30 minutes.

At the beginning of each experiment, cells were perfused with Ca^2+^-containing Ringer’s solution supplemented with 0.1% BSA. To trigger ionic responses, cells were perfused with the Caffeine (Sigma, Cat #C0750) (2mM); High concentrated potassium Ringer’s solution (High K^+^) containing: (in mM): 70 NaCl, 50 KCl, 0.8 MgCl_2_, 20 HEPES, 1.8 CaCl_2_, 15 glucose, with pH adjusted to 7.4 with NaOH; Norepinephrine (Levophed; norepinephrine bitartrate 4mg/4ml, Cat #NDC 0409-3375-04, Hospira, Inc.) (5μM), Vasopressin (Santa Cruz, Cat #sc-356188) (1μM); [Ca^2+^]c was monitored in cells loaded with Ca^2+^ specific ratiometric dye Fura2-AM (Sigma Aldrich, Cat #F0888) (1 μM), which were illuminated alternately with 340 nm and 380 nm excitations and imaged with a 510 nm long pass filter. [Ca^2+^]m was measured in cells loaded with Ca^2+^ specific dye Rhod2-AM (Biotium, Cat #50024) (1 μM) that preferentially localizes in mitochondria. Rhod2-AM was excited at 552 nm wavelength light and imaged with a 570 nm-long pass filter as previously described (Kostic, Ludtmann et al. 2015). cAMP levels were monitored in cells previously transfected/infected by using Pink Flamindo using excitation of 545 nm and imaged with a 570 nm emission filter as described before (Harada, Ito et al. 2017).

Calcium oscillations were monitored as previously described (Samanta, Mirams et al. 2018).

Bay 60-7550 (Santa Cruz, Cat #sc-396772A) and H89 (Sigma, Cat #B1427) pretreatment were performed prior to each fluorescent imaging experiment for 1h in a concentration of 1 μM and 10 μM, respectively (Monterisi, Lobo et al. 2017) (Kostic, Ludtmann et al. 2015). ATP (100 μM) (Amresco, Cat # 0220) used for mitochondrial Ca^2+^ imaging and western blot in SH-SY5Y.

### 7. Cell survival (TBOA test) and immunostaining

Triplicates of coverslips with hippocampal neurons were incubated with 80µM DL-threo-beta-benzyloxyaspartate (TBOA) (Tocris, Cat #1223) overnight as previously described (Yeh, Bulas et al. 2017). Next day, randomly chosen coverslips were incubated for 1 hour in Neurobasal-medium with or w/o Bay 60-7550 (1µM). Following the Bay 60-7550 treatment, live cells were stained with 5mM Sytox green (1:5,000) (Invitrogen, Cat #S7020) at 37^0^C and washed in Dulbesco Phospate Buffered Saline - DPBSx1 (Biological Industries, Cat# 02-020-1A).Neurons were than briefly fixed (10 minutes) with 4% paraformaldehyde (PFA, Electron microscopy sciences, Cat #BN15710) in DPBSx1, permeabilized with 0.1% TritonX-100 (Sigma Aldrich, Cat# T8787) in DPBSx1 for 2 minutes. To decrease nonspecific binding cells were immersed with 5% powdered milk (Sigma-Aldrich, Cat #70166) in DPBSx1 for 1 hour. After being washed in DPBSx1, neurons were incubated with a primary-anti NeuN (1:100) (EMD Millipore, Cat #MAB377) and secondary anti-mouse Northern Lights 557 antibody (R&D systems, Cat #NL019) containing DAPI (1: 10,000) (Sigma, Cat #D9542) respectively, each for 1 hour. Coverslips were finally mounted with Epredia™ Immu-Mount (Thermo Fisher Scientific, Cat #9990402), imaged by Nikon TiE inverted microscope driven by the NIS elements software package (Nikon) and analyzed in NIS software (described in section 6). Random 5-10 images were taken from each coverslip, neurons positively stained by NeuN were marked as areas of interest and counted manually. Counting was performed 4-10 times from each repetition in both genotypes including presence or absence of treatment. Neurons were considered live if Sytox green showed a weak or non-fluorescence, or dead if Sytox green showed strong fluorescence. The number of live cells in the field were counted blindly and averaged. The quantification represents percentage of live out of total number of cells in all images taken.

### 8. Immunoblotting

pS258-NCLX determination from whole-cell lysates, as previously described (Kostic, Katoshevski et al. 2018). Briefly, SH-SY5Y cells were first washed with DPBSx1 and lysed for 20 min in lysis buffer [Hepes 50 mM, Ph=7.4, NaCl 150 mM, EGTA 1mM, Glycerol 10%, TritonX100 1%, MgCl_2_ 10 mM, protease inhibitor (1:50) (Sigma cat # P2714) and PhosSTOP (1:10) (Roches, Cat #04 906 845 001)], all ice-cold. Lysates were then centrifuged at 4°C for 15 min at 1000*g and supernatant was collected. Equal amounts of protein (20µg) from lysates were resolved by SDS-PAGE (Molecular Biology, Cat #001981232300), transferred to the nitrocellulose membrane and Immunoblot analysis was preformed

### 9. Object recognition test

Male C57Bl/6 (Wild type-WT) mice and NCLX KO mice aged 8-14 weeks were tested for the effect of Bay 60-7550 on learning and memory using the Object Recognition (OR) test as previously described (Lueptow, Zhan et al. 2016). Each genotype group was randomly divided into 2 groups treated with either the PDE2 inhibitor Bay 60-7550 or the vehicle control. The number of mice tested is WT control (17), WT Bay 60-7550 (16), NCLX KO, vehicle and Bay 60-7550 (11). Mice were tested individually in a transparent Plexiglas box (20 × 40 cm) and filmed from above for offline analysis using Ethovision© (Noldus, The Netherlands). Behavior in the object recognition test (ORT) was assessed as previously described (Lueptow, Zhan et al. 2016). On the first day of the experiment, each mouse was placed individually in the apparatus without objects for 10 minutes to habituate to the novel environment (T0: habituation). On the second day, mice were given intraperitoneally (IP) injections of Bay 60-7550 (1 mg/kg) or vehicle [The PDE2 inhibitor Bay 60-7550 solution was dissolved in 85% saline, 10% kolliphor (Sigma, Cat #C5135-500G), 5% ethanol. The vehicle solution was dissolved in 90% saline, 10% kolliphor] 30 minutes prior to training and held in a separate cage in a different room. After 30 min, the mouse was placed in the arena in which two identical objects assembled from Lego-like blocks were placed in the test box at a distance of 16 cm from one another allowing the mouse to explore the object from all sides for 10 minutes (T1: training). Twenty-four hours later, the mice were tested in the same arena for 10 minutes with two objects: one familiar object (i.e., object used in T1) and a novel Lego object that had the same texture but differed in shape and size (T2: testing). Placement of the novel object was alternated between the two locations and counterbalanced within and between the groups. Between trials, the apparatus and each object were carefully cleaned with 80% ethanol to remove odor cues. The experimenter was blind to the treatment when testing the mice. All testing was done under low light conditions. It was expected that the mouse would explore the novel object more than the familiar object and that this difference would be greater in mice treated with Bay 60-7550, indicating that they better remembered the familiar object. Learning was assessed as the preference for the novel object over the familiar object on the third day (T3), as follows and as previously described (Lueptow, Zhan et al. 2016), briefly: Discrimination Index (DI) was calculated for each mouse during T2: [DI= (novel object exploration time - familiar object exploration time)/(novel object exploration time+familiar object exploration time)]. The data were derived from analyzing the mouse’s exploration using the Ethovision® program (Noldus, The Netherlands). A zone surrounding each object was defined by a square within a perimeter of 1 cm above the object. The time spent in the Familiar object and Novel object zone was measured for each mouse. Sitting on the object was included; however, all the films were screened to determine that no mouse spent more than 10 sec sitting on an object or engaged in other behavior that was not explorative (e.g., self-grooming, sleeping). No mice were omitted on this basis. Two mice in the WT-vehicle group, that spent less than 30 sec in the New Object zone, were eliminated from the analysis. The data were analyzed by ANOVA for the effects of genotype and drug (vehicle vs Bay 60-7550) using the Statistica ® program.

### 10. Statistical analysis

For each figure representing the fluorescence imaging experiment, at least 3 repetitions from independent cell cultures were conducted. In each of them traces were obtained from at least 3 samples of 3 regions of interest of at least 5 cells, in a non-blind manner and plotted using KaleidaGraph. All independent data points were included in the statistical analysis using 2 tailed unpaired Student’s t test after individual data point were analyzed for normality using Kolmogorov– Smirnov test. Only data that passed this latter analysis was included in the analysis for significance. No randomization or sample size calculation was performed and the study was not preregistered. The study was exploratory no exclusion of data was required and no specific test for outliers were predetermined or required. All experimental data was considered for statistical analysis. The rate of Ca^2+^ transport was derived by a linear fit of the change in the fluorescence (ΔF) over a period of 10-20 s following initiation of apparent efflux/influx (ΔF/dt) from each graph. Rates from n independent experiments (as described in legends to the figures) were averaged and displayed in bar graph format (mean ± SEM). GraphPad Prism 6 software was used for statistical analysis and Adobe Illustrator for the graphic design of the figures. All bar graphs were presented as averaged single sample values of (n) independent measurements ± SEM. A value of p < 0.05 was accepted as statistically significant. Symbols of significance used: ns p > 0.05, * p ≤ 0.05, ** p ≤ 0.01, *** p ≤ 0.001, **** p ≤ 0.0001.

## Acknowledgements

This study was funded by (ISF 1424/17, DIP SE 2372/1-1 to Israel Sekler)

## Author contribution

Maya Rozenfeld, Ivana Savic Azoulay, Israel Sekler wrote the manuscript; Maya Rozenfeld, Ivana Savic Azoulay, Alexandra Stavsky and Tsipi Ben Kasus Nissim performed the experiments; Michal Hersfinkel, Ora Kofman, Israel Sekler provided experimental oversight and wrote the paper; Maya Rozenfeld, Ivana Savic Azoulay, Ora Kofman and Israel Sekler designed the experiments; Maya Rozenfeld,, Ivana Savic Azoulay, Israel Sekler and Ora Kofman interpreted the data.

---

**Supplementary Figure 1.**
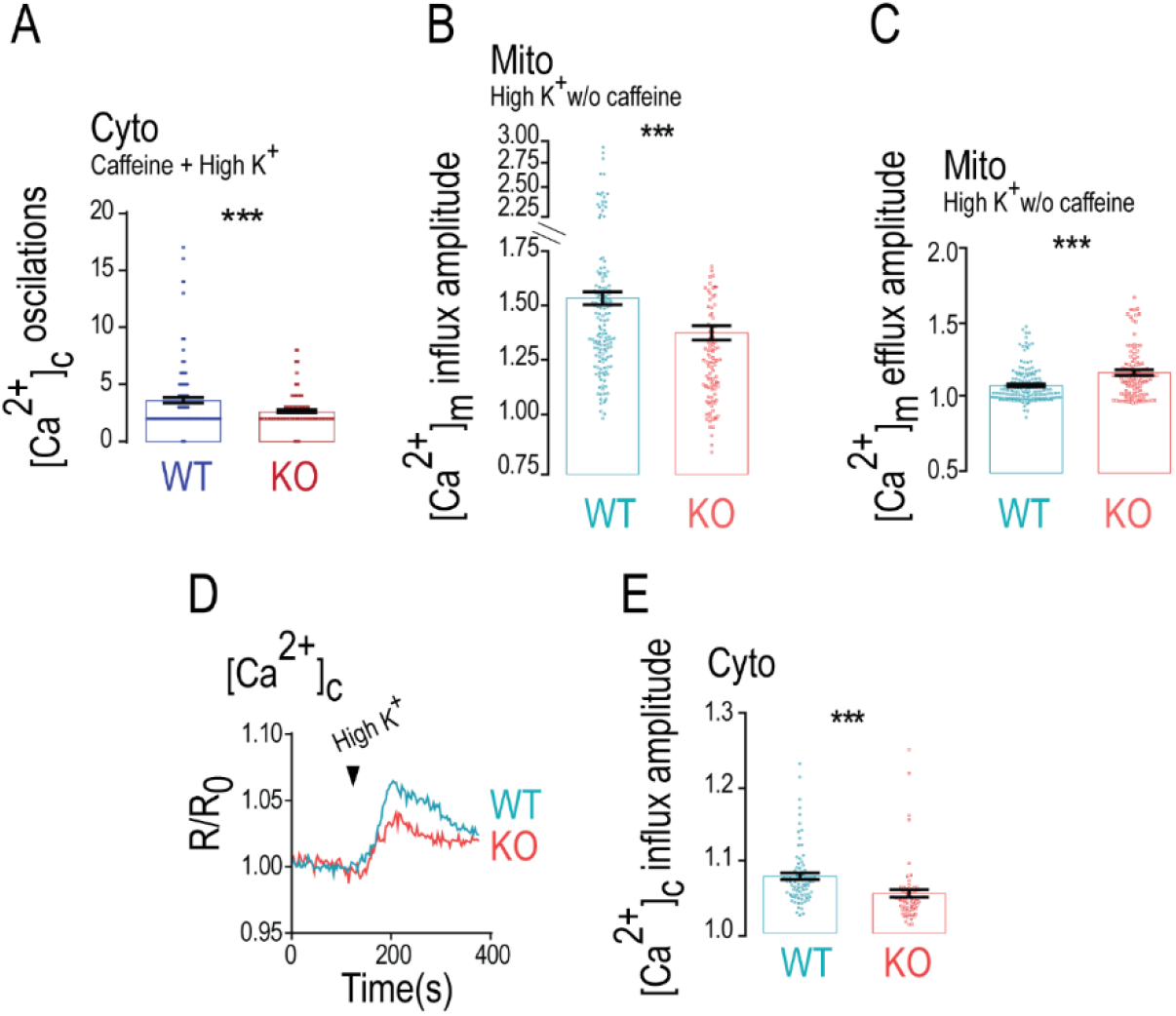
Caffeine effects on [Ca^2+^]c and [Ca^2+^]m amplitudes are NCLX dependent. (A) Quantification of [Ca^2+^]c oscillations showed in (Figure 1G) for WT (n=170) and NCLX KO neurons (n=166); (B) Quantification of [Ca^2+^]m influx amplitude showed in (Figure 1I) for WT (n=160) and NCLX KO neurons (n=114); (C) Quantification of [Ca^2+^]m efflux amplitude showed in (Figure 1I) for WT (n=160) and NCLX KO neurons (n=114); (D) Representative fluorescence traces of [Ca^2+^]c changes in WT and NCLX KO neurons triggered by high K^+^ Ringer’s solution; (E) Quantification of [Ca^2+^]c influx amplitude changes during the maximum phases showed in (D) for WT (n=91) and NCLX KO neurons (n=63); All summary data represent mean ± SEM., *** p < 0.001.

**Supplementary Figure 2.**
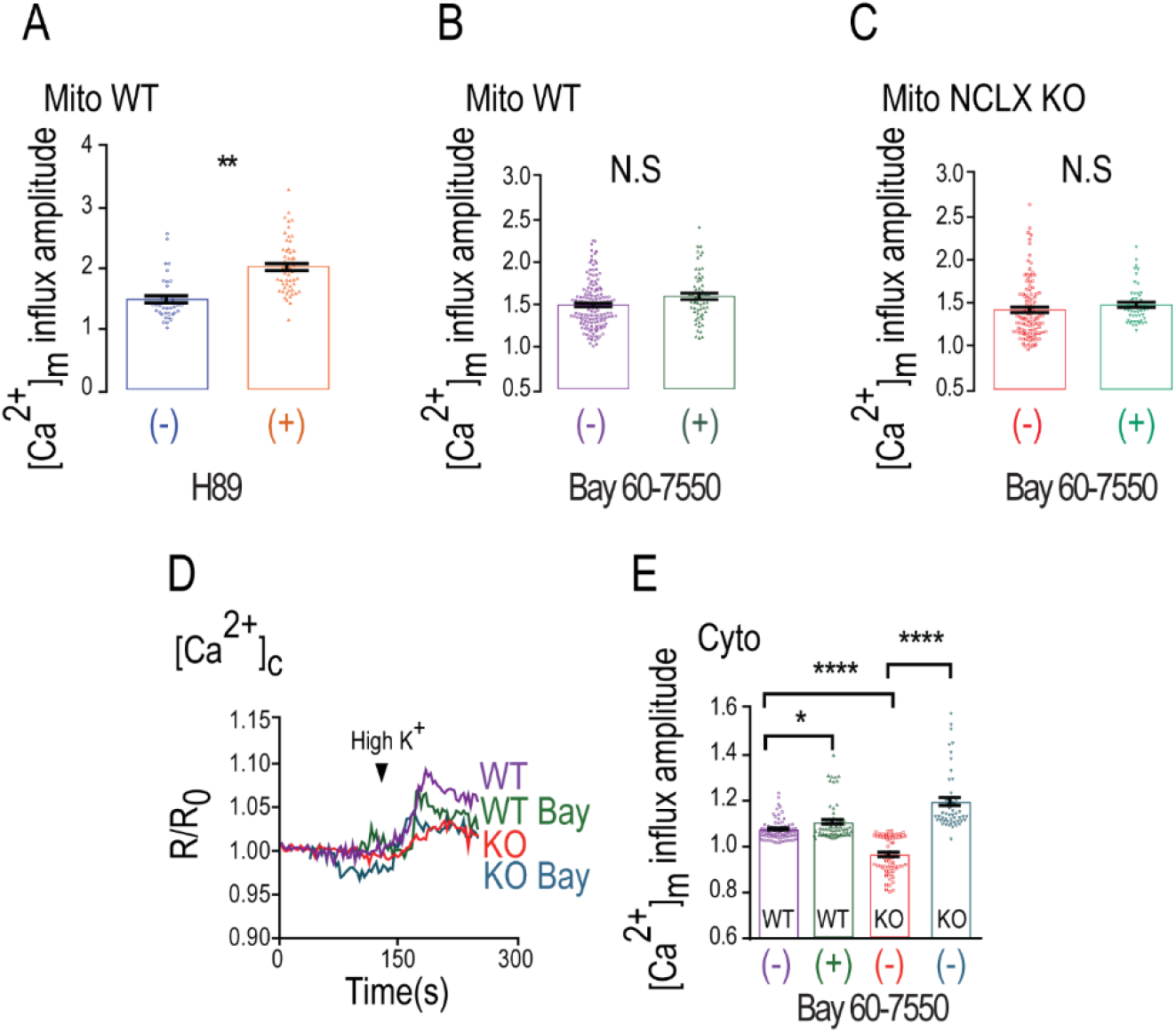
Bay 60-7550 does not affect PKA dependent-[Ca^2+^]m and [Ca^2+^]c amplitudes neither in WT nor NCLX KO neurons. (A) Quantification of [Ca^2+^]m influx amplitude showed in (Figure 2A) for control (n=38) and H89 treated WT neurons (n=54); (B) Quantification of [Ca^2+^]m influx amplitude showed in (Figure 2G) for control (n=140) and Bay 60-7550 treated WT neurons (n=60); (C) Quantification of [Ca^2+^]m influx amplitude changes showed in (Figure 2J) for control NCLX KO (n=100) and Bay 60-7550 treated NCLX KO neurons (n=52); (D) Representative fluorescence traces of [Ca^2+^]c monitored in control and Bay 60-7550 pretreated WT and NCLX KO neurons; (E) Quantification of [Ca^2+^]c influx amplitude changes showed in (S2D) for control (n=91) and Bay 60-7550 treated (n=72) WT, control (n=63) and Bay 60-7550 pretreated (n=47) NCLX KO neurons; All summary data represent mean ± SEM., * p < 0.05, ** p < 0.01**** p < 0.0001, N.S- non-significant.

**Supplementary Figure 3.**
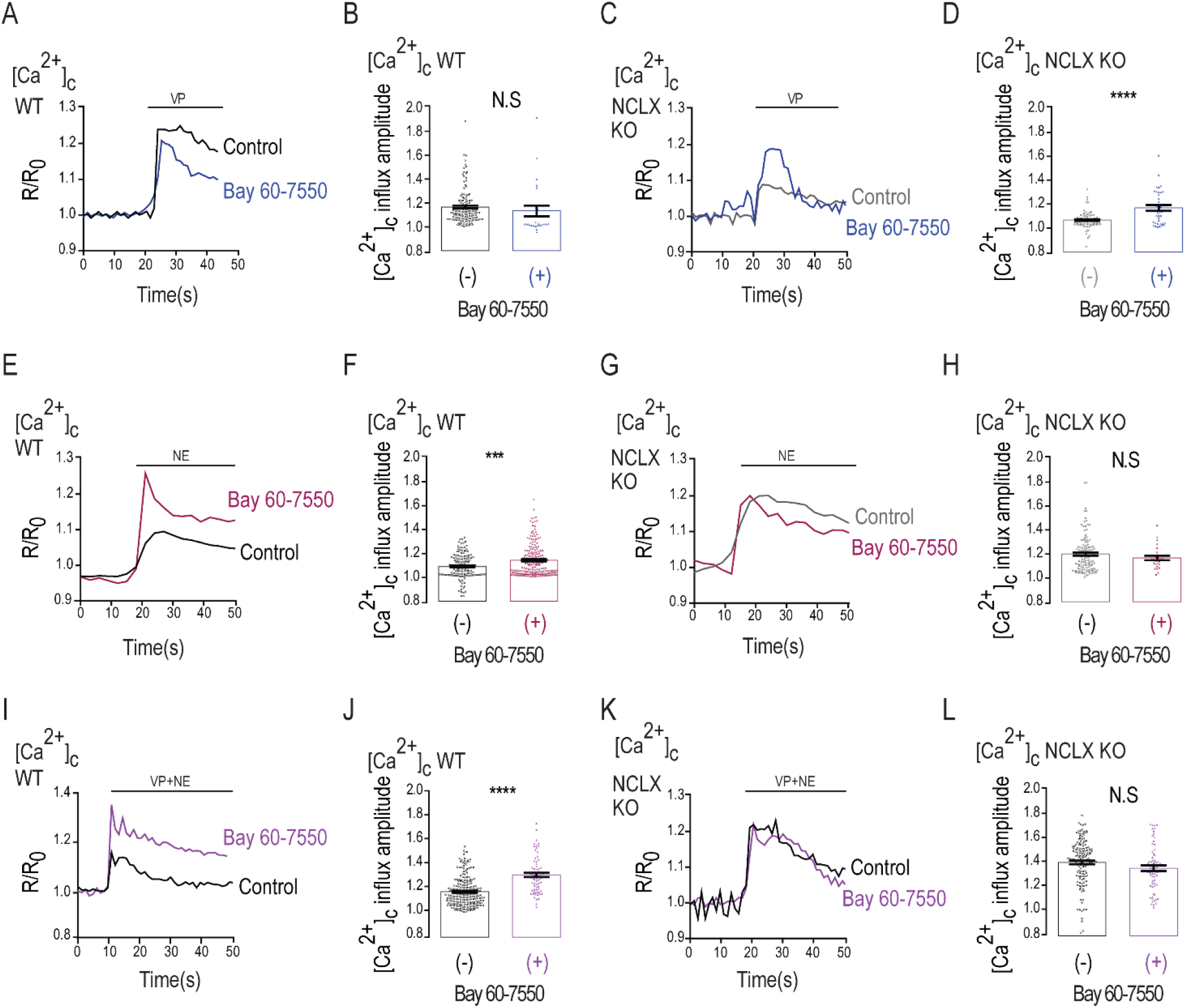
VP and NE triggered [Ca^2+^]c signals are not PDE2 dependent. (A) Representative fluorescence traces of [Ca^2+^]c monitored in control and Bay 60-7550 pretreated WT neurons. Neurons were preloaded with Fura2-AM (1 µM), initially superfused with Ringer’s solution and then by VP containing Ringer’s solution; (B) Quantification of [Ca^2+^]c influx amplitudes showed in (A) for control (n=110) and Bay 60-7550 pretreated WT neurons (n=28); (C) Representative fluorescence traces of [Ca^2+^]c changes in control NCLX KO and Bay 60-7550 pretreated NCLX KO neurons of the same experiment done as in (A); (D) Quantification of [Ca^2+^]m influx amplitudes showed in (C) for control NCLX KO (n=70) and Bay 60-7550 pretreated NCLX KO neurons (n=40); (E) Representative fluorescence traces of [Ca^2+^]c changes monitored in control and Bay 60-7550 pretreated WT neurons of the experiment done as in (A), here [Ca^2+^]c was triggered by NE containing Ringer’s solution; (F) Quantification of [Ca^2+^]c influx amplitudes showed in (E) for control (n=140) and Bay 60-7550 pretreated WT neurons (n=190); (G) Representative fluorescence traces of [Ca^2+^]c changes in control NCLX KO and Bay 60-7550 pretreated NCLX KO neurons of the same experiment done as in (E); (H) Quantification of [Ca^2+^]c influx amplitudes rates showed in (G) for control NCLX KO (n=122) and Bay 60-7550 pretreated NCLX KO neurons (n=27); (I) Representative fluorescence traces of [Ca^2+^]c changes monitored in control and Bay 60-7550 pretreated WT neurons of the experiment performed as in (A), with [Ca^2+^]c signal evoked by mix of VP and NE of the same concentrations as in (A and E); (J) Quantification of [Ca^2+^]c efflux rates showed in (I) for control (n=218) and Bay 60-7550 pretreated WT neurons (n=68); (K) Representative fluorescence traces of [Ca^2+^]c monitored in control NCLX KO and Bay 60-7550 pretreated NCLX KO neurons of the experiment performed as in (I); (L) Quantification of [Ca^2+^]c efflux rates showed in (K) for control NCLX KO (n=149) and Bay 60-7550 pretreated NCLX KO neurons (n=68); All summary data represent mean ± SEM., *** p < 0.001 **** p < 0.0001, N.S- non-significant.

